# Polycomb Repressive Complex 1 subunit Cbx4 positively regulates effector responses in CD8 T cells

**DOI:** 10.1101/2022.10.03.510675

**Authors:** G.A. Melo, T. Xu, C. Calôba, A. Schutte, G. Brum, T.O. Passos, L. Higa, A. Gonçalves, A. Tanuri, J.P.B. Viola, M.B.F. Werneck, H.I. Nakaya, M.E. Pipkin, G.J. Martinez, R.M Pereira

**Affiliations:** Departamento de Imunologia, Instituto de Microbiologia Paulo de Góes, Universidade Federal do Rio de Janeiro, 21941-902, Rio de Janeiro, RJ, Brazil; Center for Cancer Cell Biology, Immunology, and Infection; Discipline of Microbiology and Immunology. Rosalind Franklin University of Medicine and Science, 3333 Green Bay Road, North Chicago, IL 60064, USA; Department of Immunology and Microbiology, UF Scripps Biomedical Research, University of Florida, Jupiter, FL 33458, USA; Hospital Israelita Albert Einstein, 05652-900, São Paulo, SP, Brazil; Instituto de Biofísica Carlos Chagas Filho, Universidade Federal do Rio de Janeiro, 21941-902, Rio de Janeiro, RJ, Brazil; Programa de Imunologia e Biologia Tumoral, Instituto Nacional do Câncer, 20231-050, Rio de Janeiro, RJ, Brazil; Departamento de Genética. Instituto de Biologia, Universidade Federal do Rio de Janeiro, 21941-902, Rio de Janeiro, RJ, Brazil; Department of Clinical and Toxicological Analyses, School of Pharmaceutical Sciences, University of São Paulo, 05508-000, São Paulo, SP, Brazil

**Keywords:** Epigenetics, Polycomb, Cbx4, PRC1, T cell differentiation, CD8 T cell

## Abstract

CD8 T cell differentiation is controlled by the crosstalk of various transcription factors and epigenetic modulators. Uncovering the different players in regulating this process is fundamental to improving immunotherapy and designing novel therapeutic approaches. Here, we show that Polycomb Repressive Complex (PRC)1 subunit Chromobox (Cbx)4 favors differentiation to effector CD8 T cells. Cbx4 deficiency in CD8 T cells induced transcriptional signature and phenotype of memory cells, increasing the formation of memory population during acute viral infection. It has been previously shown that besides chromodomain-mediated binding to H3K27me3, Cbx4 function as a SUMO E3 ligase in a SUMO interacting motifs (SIM)-dependent way. The overexpression of Cbx4 mutants in distinct domains showed that this protein regulates CTL differentiation primarily in a SIM-dependent way and partially through its chromodomain. Our data revealed a novel role of a Polycomb group protein Cbx4 controlling CD8 T lymphocyte differentiation and indicates the SUMOylation process as a key molecular mechanism connected to chromatin modification in this process.

**Summary:** Understanding the epigenetic control of CTL differentiation is critical for the manipulation of these cells in immunotherapy protocols. This article demonstrates a novel role for Cbx4, a Polycomb-group protein, in supporting CD8 T cell commitment to an effector cell phenotype.

## Introduction

CD8 T cells are an essential component of adaptive immunity, working as key players in the elimination of intracellular pathogens and tumor cells (1). Upon activation, CD8 T cells undergo intense expansion and differentiate into a heterogeneous population composed of pro-inflammatory and cytotoxic terminal effector cells (TE), characterized by expression of the killer cell lectin-like receptor G1 (KLRG1^+^) and low expression of IL-7Ra (CD127^-^), and memory precursor (MP) cells that are responsible for the generation of long-term immunity, that express CD127 and low levels of KLRG1 (KLRG1^-^CD127^+^). Complex transcriptional programs that are regulated dynamically control gene expression that specifies the differentiation of these distinct CD8 T cell states. The identification of the key players that participate in this process is critical for the understanding of these molecular processes and for advancing novel therapeutic strategies (1, 2). Several transcription factors have been described to drive CD8 T cell differentiation, including T-bet (3, 4), Blimp1 (5, 6), Zeb-2 (7, 8), Id2 (9–11), and NFAT1 (12) which promote generation of TE cells, and Tcf-1 (13, 14), Eomes (3, 15), Bcl-6 (16, 17), Id3 (18, 19), Foxo1 (20–22) and Runx3 (23) for memory cell differentiation. Transcription factors cooperate with epigenetic regulators, including chromatin modifiers that add, remove or recognize post-translational modifications in histones, adding a new level of complexity to this differentiation process that remains to be completely uncovered (24–26).

The Polycomb Repressor Complex (PRC)2, responsible for tri-methylating lysine 27 of histone 3 (H3K27me3), which is considered a repressive mark, was described as having an essential role in commitment to effector cell differentiation through the deposition of this mark in memory-related genes, leading to transcriptional repression during acute viral infection (25, 27, 28). In addition, members of this complex were upregulated right after activation in cells that later differentiated into terminal effector CD8 T cells (28). Furthermore, it was demonstrated that H3K27me3 removal by lysine demethylase Kdm6 promotes effector differentiation through de-repression of effector genes (29–31). These evidences underscore the importance of H3K27me3 methylation/demethylation dynamic regulation to CD8 T cell differentiation. H3K27me3 can recruit another epigenetic complex from the Polycomb group (PcG), the PRC1, that also works as a repressor through the ubiquitylation of lysine 119 of histone H2A (H2AK119ub) by its catalytic subunit Ring1, as well as through chromatin remodeling mechanisms mediated by other subunits that can integrate the complex (32). PRC1 composition is highly heterogeneous, and its distinct configurations are divided in canonical (cPRC1) or noncanonical PRC1 (ncPRC1) (33, 34). While ncPRC1 complexes can be recruited to genome sites independently of PRC2 function (35, 36), cPRC1 are known to be recruited to H3K27me3-tagged *loci*. This process is thought to be mediated by recognition and binding of Chromobox (Cbx) family proteins (Cbx2, Cbx4, Cbx6, Cbx7, and Cbx8) to this repressive epigenetic mark through its conserved chromodomain (32, 37). Cbx4 is reported to function additionally as a small ubiquitin-like modifier (SUMO) E3 ligase through its two SUMO interacting motifs (SIM) (38, 39). SUMO conjugation is a dynamic post-translational modification that can alter the molecular interactions, stability and activity of targeted proteins and mediates the regulation of several critical cellular processes, such as control of gene expression and chromatin structure (40, 41). Besides, Cbx4 E3 SUMO ligase activity has been reported to regulate essential transcriptional regulators, such as Dnmt3a and Hif-1α (42, 43).

Here we show that Cbx4 induces effector cell commitment primarily through its SIM domain-related mechanisms. Cbx4-deficient CD8 T cells showed increased expression of memory-related surface markers CD127 and CD62L for both *in vitro* differentiation and during acute viral infection. Consistent with our phenotyping results, RNA-Seq of *in vivo*-differentiated Cbx4-deficient cells showed skewing towards memory precursor transcriptional signature. Furthermore, Cbx4 deficiency led to increased memory generation at acute lymphocytic choriomeningitis virus (LCMV) viral infection memory time point. To rescue the Cbx4 deficient phenotype, we overexpressed Cbx4 or Cbx4 functional domain mutants in CD8 T cells. We found that the overexpression restored the effector population in a SIM domain-dependent manner and only partially depended on its chromodomain. Our data, suggest that Cbx4 promotes CD8 T cell effector differentiation, which is mainly driven by SUMOylation mechanisms and less so by its capacity to bind to methylated histones.

## Methodology

### Mice

All mice were on a C57BL/6 background. The experimental mice were 6-to 8-wk-old and sex- and age-matched. P14 (LCMV gp33-41-H2-Db–specific) Thy1.1 and CD4-Cre mice have been previously described (12). Cbx4^fl/fl^ mice (obtained from Dr. Guo-Liang Xu, Institute of Biochemistry and Cell Biology, China (44)) were bred to create the following genotypes: Cbx4^fl/fl^Cd4^Cre^ and P14 Thy1.1^+^ Cbx4^fl/fl^Cd4^Cre^. Mice were housed in specific pathogen-free barrier facilities and used according to protocols approved by the Rosalind Franklin University of Medicine and Science Institutional Animal Care and Use Committee (IACUC). Mice housed in the Mouse facility of Departamento de Imunologia (UFRJ) were used according to the rules established by CONCEA (Conselho Nacional de Experimentação Animal) and UFRJ CEUA (Comitê de Ética no Uso de Animais) approval (Protocol 054/20).

### T cell isolation and culture

Naive CD8^+^ T cells were purified from spleen and lymph nodes using the Mouse Naive CD8^+^ T Cell Isolation Kit (StemCell Technologies) or through negative selection using Dynabeads Untouched Mouse CD8 cells Kit (Thermo Fisher Scientific) followed by Fluorescence-Activated Cell Sorting (FACS). Naïve CD8 T cells were cultured on Dulbecco’s modified Eagle’s Medium (DMEM) supplemented with 10% heat-inactivated FBS, 2 mM L-glutamine, 100 U/mL penicillin-streptomycin, 50 ng/ml gentamicin, 1X MEM non-essential amino acids, 1 mM sodium pyruvate, 1X MEM Vitamin Solution, 10 mM HEPES, and 50 μM β-Mercaptoethanol (45). Naïve CD8 T cells were activated with 1 μg/ml (effector-like polarization) or 50 ng/mL (memory-like polarization) of anti-CD3 (clone 2C11) and 1 μg/mL of anti-CD28 (clone 37.51) (Thermo Fisher Scientific) at 1 million cells per mL on a 12-or 24-well plate that had been precoated with 300 μg/ml goat anti-hamster IgG (Pierce Protein Biology, Life Technologies) (45). Cells were removed from the initial stimulus 48 h after activation and were cultured at 0.5 million/mL in the presence of 100 U/ml of recombinant human (rh)IL-2 for effector-like or 10 U/mL of rhIL-2, 10 ng/mL of rhIL-7 and 10 ng/mL of rhIL-15 for memory-like phenotype polarization. In indicated experiments, 200 U/mL of recombinant murine (rm)IL-2 were used to generate effector-like cells and 20 U/mL of rhIL-2, 10 ng/mL of rmIL-7 and 10 ng/mL of rmIL-15 for memory-like cells (46).

### Retroviral plasmids and transduction

For Cbx4 knockdown experiments, retroviral particles were generated by transfecting Phoenix-Eco cells with Ametrine-expressing murine retroviral vectors containing shRNAs targeting CD4 (shCD4) or Cbx4 (shCbx4) mRNA (47). For Cbx4 over-expression, Plat-E cells were transfected with GFP-expressing murine retroviral vectors containing unaltered Cbx4 coding sequence (Cbx4-OE), Cbx4 mutants carrying two point mutations (F11A/W35L) at Cbx4 chromodomain (ΔChromo), deletion of the two SUMO interacting motifs (SIM) domains (ΔSIM1-2) or GFP-expressing empty vector (Mock), kindly donated by Dr. Wang (48). Virus supernatant was filtered through 0.45-mm filters and used fresh for transduction in Cbx4 knockdown experiments or concentrated by centrifugation at 6000 x g (F14-14 × 50cy rotor) at 4ºC overnight for Cbx4 over-expression experiments. *In vitro*-activated CD8 T cells (as described above) were transduced with retroviral particles 18-20h after activation. T cell culture medium was carefully replaced with media containing fresh or concentrated retrovirus supplied with 8 μg/ml polybrene and centrifuged at 800 x g for 1 h at 37ºC and then put into a 37ºC 10% CO2 incubator for 4h. For the adoptive transfer of transduced P14 cells, T cell cultures were immediately harvested after incubation. For *in vitro* culture experiments, the conditioned T cell media was replaced, and cells were then expanded until day 6 or day 14 for effector-or memory-like polarization, respectively.

### LCMV infection and adoptive cell transfer

Mice were infected i.p. with 2 × 10^5^ PFUs of LCMV Armstrong (LCMV Arm) and analyzed on day 45 (d45) post-infection (p.i.). For adoptive transfer experiments with transduced CD8 T cells, congenic C57BL/6 (Thy1.2) mice received intravenously 5×10^5^ *in vitro*-transduced P14 Thy1.1 cells (expressing shCD4 or shCbx4) and were subsequently infected i.p. with 1.5 × 10^5^ PFUs of LCMV clone13, as previously described (47). LCMV strains were initially provided by Dr. Shane Crotty, La Jolla Institute, CA and expanded with BHK cells as described before (49).

### Flow Cytometry analysis

Spleens and lymph nodes were used for isolating single-cell suspension. RBCs were lysed from spleens with ACK lysis buffer. For LCMV tetramer staining, H2Db-gp33-41 (KAVYNFATC) Alexa Fluor 647 or APC was incubated at room temperature before staining for cell surface molecules and intracellular staining. Cytokine production was measured by FACS from *ex vivo* splenic cells or *in vitro* cultured cells upon restimulation with 10 nM PMA and 1 μM Ionomycin for 4 h in the presence of brefeldin A. Samples were run on FACSAria IIu or LSRII (BD Biosciences), and data were analyzed with FlowJo (Version 9.9.4 and Version 10.7.1).

### Cytotoxicity assay

For cytotoxicity assay, naïve P14 CD8 T cells were activated *in vitro*, transduced with shCD4 or shCbx4, and cultured with 10 U/ml rhIL-2 as described above. On day 6, Ametrine^+^ cells were purified by FACS and co-cultured at different ratios with GFP-expressing parental mammary carcinoma cell line EO771 (negative control to determine nonspecific target lysis) or EO771 cells expressing the cognate Ag gp33-41 (EO771-GFP-gp33-41). After overnight incubation (12 h), the remaining live GFP-expressing EO771 cells were determined by flow cytometry as a measure of the cytotoxic activity. EO771 cells cultured in the absence of CTLs were used as a baseline for cell death.

### RNA sequencing

FACS-purified cells from *in vivo* experiment were washed twice with PBS and used for RNA sequencing. RNA-seq libraries were prepared using SMARTer Stranded RNA-Seq Kit (Clontech) according to the manufacturer’s instructions. RNA-Seq libraries were sequenced with the rapid run protocol with a HiSeq 2500 instrument (Illumina). The paired-end reads that passed Illumina filters were filtered for reads aligning to tRNA, rRNA, adapter sequences, and spike-in controls. The reads were then aligned to the UCSC mm9 reference genome using TopHat (v1.4.1). DUST scores were calculated with PRINSEQ Lite (v 0.20.3), and low complexity reads (DUST > 4) were removed from the BAM files. Read counts to each genomic feature were obtained with htseq-count (v 0.6.0) using the default “union” option. After removing absent features (zero counts in all samples), the raw counts were imported to R/Bioconductor package DESeq2 (v1.24.0) to identify differentially expressed genes among samples. We considered genes differentially expressed between two groups of samples when the DESeq2 analysis resulted in a p-value < 0.05. Gene set enrichment analysis (GSEA), that considers the gene expression ranking to calculate enrichment, was performed by comparing the differentially expressed genes between shCD4 and shCbx4-treated samples (pre-ranked gene list) to published SLEC and MPEC datasets (gene set) (23).

### Quantitative real-time RT-PCR

Total RNA was isolated from FACS-purified CD8 T cells using TRIzol reagent (Invitrogen) according to manufacturer’s instructions. Superscript reverse transcriptase (Invitrogen) and oligonucleotide primers were used to synthesize cDNA. Gene expression was examined with 7500 Real-Time PCR System (Applied Biosystems) using Power SYBR green PCR Master Mix (Thermo Fisher Scientific). Gene expression was normalized to *Rpl22* (encodes L22 ribosomal protein) gene expression. The following primers were used: *Rpl22* forward, 5’-ACCCTGGACTGCACTCACCCTG-3’; *Rpl22* reverse, 5’-CCGCCGAGGTTGCCAGCTTT-3’; Sell forward, 5’-CATTCCTGTAGCCGTCATGG-3’; Sell reverse, 5’-AGGAGGAGCTGTTGGTCATG-3’; Il7r forward, 5’-GCGGACGATCACTCCTTCTG-3’; Il7r reverse, 5’-AGCCCCACATATTTGAAATTCCA-3’; *Cbx4* forward, 5’-AGTGGAGTATCTGGTGAAATGGA-3’; *Cbx4* reverse, 5’-TCCTGCCTTTCCCTGTTCTG-3’.

### Statistics and analysis

Graphs were plotted using Prism 7 or 8 (GraphPad). Statistical analysis was performed using nonpaired one-way ANOVA followed by Tukey multiple comparisons, two-tail paired or nonpaired Student t test, or two-way ANOVA followed by Tukey, Sidak or Dunnet multiple comparisons, as indicated.

## Results and Discussion

### Cbx4 deficiency skews CD8 T cell differentiation to a memory phenotype and impacts CD8 T cell cytotoxicity

To assess the role of Cbx4 in CD8 T cell differentiation, we employed LCMV acute infection model using adoptive transfer of CD8 T cells from P14 mice. Naïve CD8 T cells from P14 Thy1.1^+^ mice were activated *in vitro* and transduced with retroviral vectors (RV) expressing shRNA targeting Cbx4 mRNA (shCbx4) or CD4 mRNA as a control (shCD4). Transduced cells were then adoptively transferred to WT Thy1.2^+^ congenic mice that were infected on the same day with LCMV, as previously described (47) (Fig.1.A). The efficiency of shRNA-mediated silencing of Cbx4 was confirmed by RT-qPCR (Suppl.Fig1. A).

**Figure 1.**
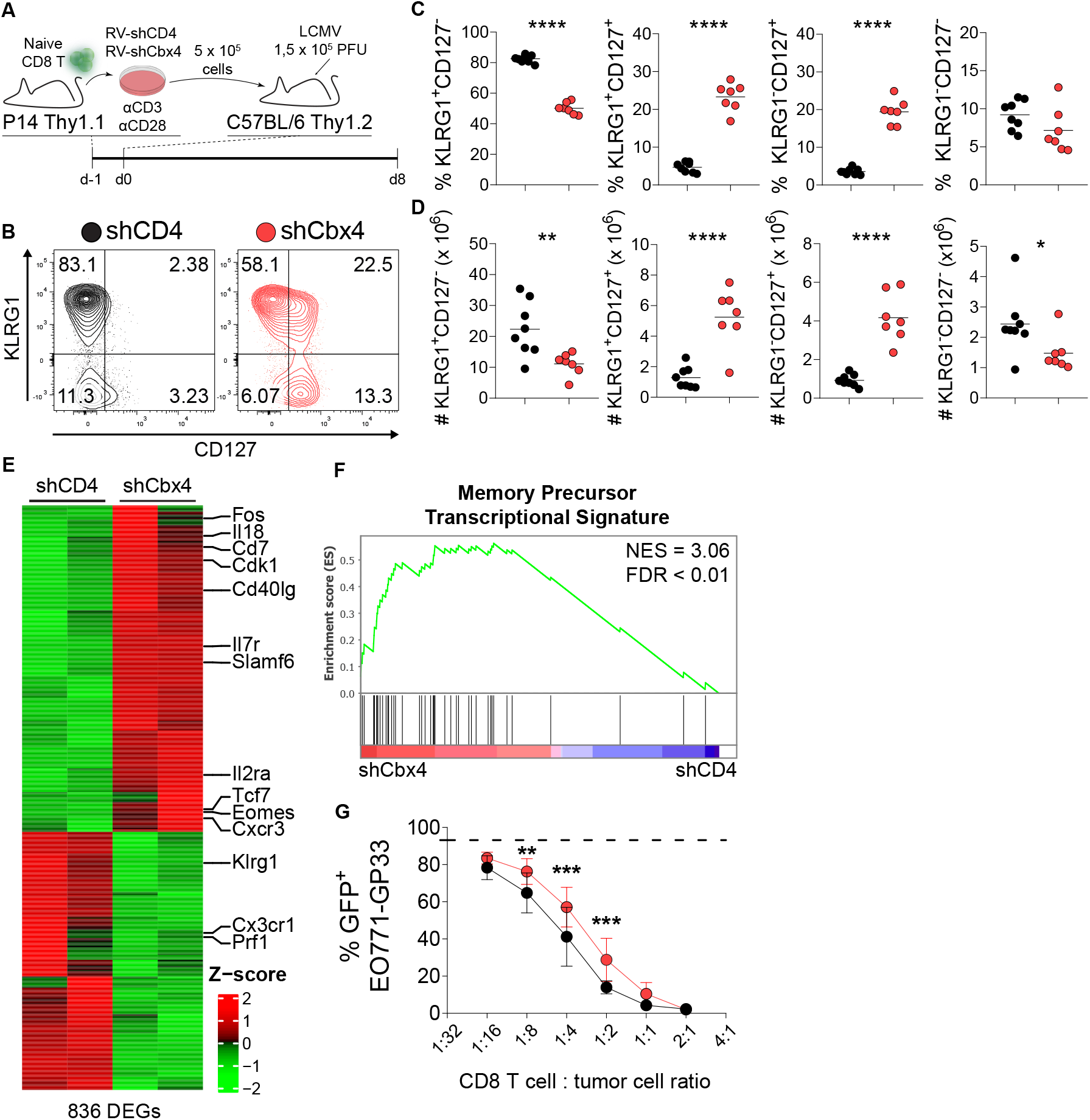
Cbx4 deficiency skews CD8 T cell differentiation to a memory phenotype. (A) *In vitro*-activated P14 Thy1.1^+^ cells transduced with retrovirus (RV) expressing control shRNA (shCD4, black) or shRNA targeting Cbx4 (shCbx4, red) were transferred to congenic receptor mice subsequently infected with LCMV and analyzed by flow cytometry for KLRG1 and CD127 expression in Ametrine^+^ Thy1.1^+^ CD8^+^ T cells at 8 dpi (B) or Ametrine^+^-sorted for RNA-Seq. Summary of frequency (C) and total cell numbers (D) of populations analyzed in KLRG1 x CD127 gate. RNA-Seq results showed 836 differentially expressed genes (DEGs) that were visualized in a heatmap of Z-score values clustered by hierarchical clustering (E). DEGs were tested by GSEA for enrichment of transcription signature from memory precursor (KLRG1^lo^CD127^hi^) population from day 8 of acute LCMV infection (23) (F). P14 shCD4 and shCbx4 cells were also assessed for *in vitro* specific cytotoxicity (dashed line indicates baseline tumor cell survival) (G). Representative contour plots for KLRG1 and CD127 expression are shown (B). Data are representative of two independent experiments (n ≥ 3). \**p* < 0.05, \*\**p* < 0.01, \*\*\**p* < 0.001, \*\*\*\**p* < 0.0001 by unpaired two-tailed Student *t* test (C, D).

The differentiation of P14 cells to effector or memory phenotype was analyzed 8 days post-infection by measuring KLRG1 and CD127 expression (Fig.1.B-D). Transferred LCMV-specific Cbx4-deficient CD8 T (P14 shCbx4) cells had lower frequency and number of TEs (KLRG1^+^CD127^-^) and, accordingly, higher frequency and cell number of populations expressing the memory-associated marker CD127, both in MP (KLRG1^-^CD127^+^) and in KLRG1^+^CD127^+^ double positive population. The frequency of transferred Thy1.1^+^ P14 cells and transduced Ametrine^hi^ Thy1.1^+^ P14 cells were lower in mice receiving P14 shCbx4 cells in comparison to mice receiving control cells; however, no significant difference was observed in the cell numbers, which is consistent with the increase in total CD8^+^ T cell population numbers in mice receiving P14 shCbx4 cells (Suppl.Fig.1. B).

To further investigate the contribution of Cbx4 to the regulation of CD8 T cell transcriptional profile, P14 CD8 T cells from the LCMV acute infection model described above were sort-purified on d8 post-infection, and total RNA was extracted for RNA-Seq. Data analysis showed 836 differentially expressed genes (DEGs) when comparing P14 shCbx4 CD8 T cells to P14 shCD4 CD8 T cells (Fig.1.E). Among those genes, several surface markers that positively correlate with the memory cells were upregulated on Cbx4 deficient cells, including *Il7r* (CD127) and *Slamf6* (2, 50). In addition, genes encoding key transcription factors *Eomes* and *Tcf7* (Tcf-1), which promote the generation and persistence of central memory CD8 T cells, were also upregulated in comparison to control cells (3, 13–15). Concordantly, genes positively correlated with the effector phenotype and function, such as *Klrg1, Prf1* and *Cx3cr1*, were downregulated in P14 shCbx4 compared to shCD4 CD8 T cells. These gene expression alterations, such as the upregulation of *Tcf7* and *Il7r*, were not just due to a higher frequency of memory cells in our Cbx4-deficient population, as sorted Cbx4-deficient KLRG1^hi^ CD8 T cells also showed an enrichment of memory-associated genes (Suppl.Fig.1. D). To test whether these gene expression alterations lead to a global shift towards memory transcriptional programs, we performed Gene Set Enrichment Analysis (GSEA) using previously published memory precursor (KLRG1^lo^CD127^hi^) transcriptional signature from day 8 acute LCMV infection model (23). We found that genes upregulated in P14 shCbx4 cells showed enrichment in memory precursor transcriptional signature (Fig.1.F), indicating that Cbx4 could directly regulate memory-associated genes. It is also plausible that Cbx4 could indirectly impact the expression of transcription factors related to CD8 T cell differentiation by controlling other transcriptional regulators. For example, our RNA-Seq showed that *c-Myb*, a transcriptional activator of *Tcf7* (51), is upregulated in P14 shCbx4 cells. Similarly, Cbx4 was reported to activate the Wnt/β-catenin pathway, a pathway upstream of TCF-1 activation, in human lung adenocarcinoma cells (52). Future studies will be focused on determining which transcriptional regulators involved in CD8 T cell differentiation are directly regulated by Cbx4.

As we observed the downregulation of genes related to effector function (i.e.: *Prf1*), we explored if Cbx4 deficiency could impact CD8 T cell cytotoxic function. P14 cells were analyzed in an *in vitro* cytotoxicity assay, co-culturing activated and retrovirally transduced P14 cells with GP33-expressing GFP^+^ EO771 tumor cells (EO771) (Fig.1.G). We observed that P14 shCbx4 cells had diminished cytotoxicity compared to P14 shCD4, which further supports the observation of defective effector differentiation with concomitant skewing to memory phenotype in the absence of Cbx4 protein.

Collectively, our findings shows that Cbx4 deficiency upregulates memory precursor transcriptional signature and expression of memory surface markers (CD127, CD62L), and decreases effector cytotoxic function, revealing a skewing of CD8 T cells towards a memory phenotype and a role for the Polycomb protein Cbx4 in the control of CD8 T cell differentiation.

### Cbx4 deficiency leads to increased memory CD8 T cell formation

To confirm if Cbx4 could control memory CD8 T cell formation, we used the acute LCMV infection in T cell-specific Cbx4 deficient mice (Cbx4^fl/fl^Cd4^Cre^, herein referred to as Cbx4 TKO mice), analyzing polyclonal Cbx4 deficient CD8 T cells 60 days post-infection (Fig.2.A). Analysis of lymphocyte populations on thymus, lymph nodes and spleens of Cbx4 T KO mice at steady state showed no alteration in T cell development or abnormal phenotype of lymphocytes in secondary lymphoid organs (data not shown). On day 60 post-infection, Cbx4 TKO mice showed a slight increase in the frequency and number of CD8 T cells, as well as higher number of LCMV-specific H2Db-gp33-41^+^ (GP33^+^) cells (Fig.2.B, C). In addition, Cbx4 TKO mice had higher frequency and cell number of KLRG1^+^CD127^+^ population in GP33^+^ CD8 T cells (Fig.2. D-F). We also observed higher number of memory KLRG1^-^CD127^+^ population, although this was not reflected in population frequency. Accordingly, we found decreased frequency and cell number of terminal effector (KLRG1^+^CD127^-^) GP33^+^ CD8 T cells (Fig.2.D-F). The expression of CXCR3, which is associated with memory population formation and rapid recall response (53), was upregulated on GP33^+^ CD8 T cells of Cbx4 TKO mice, as seen by the increased cell numbers of KLRG1^+^CXCR3^+^ and KLRG1^-^CXCR3^+^ populations (Suppl.Fig.2.A, B). To further validate the enrichment of memory-associated genes in Cbx4 deficient CD8 T cells, we performed intracellular staining to examine expression of transcription factor T-bet and Eomes from our *ex vivo* samples. We observed that T-bet/Eomes ratio was significantly lower in Cbx4 TKO mice, regardless of KLRG1 expression, in line with a global increased memory profile upon Cbx4 deficiency (Fig.2.G).

**Figure 2.**
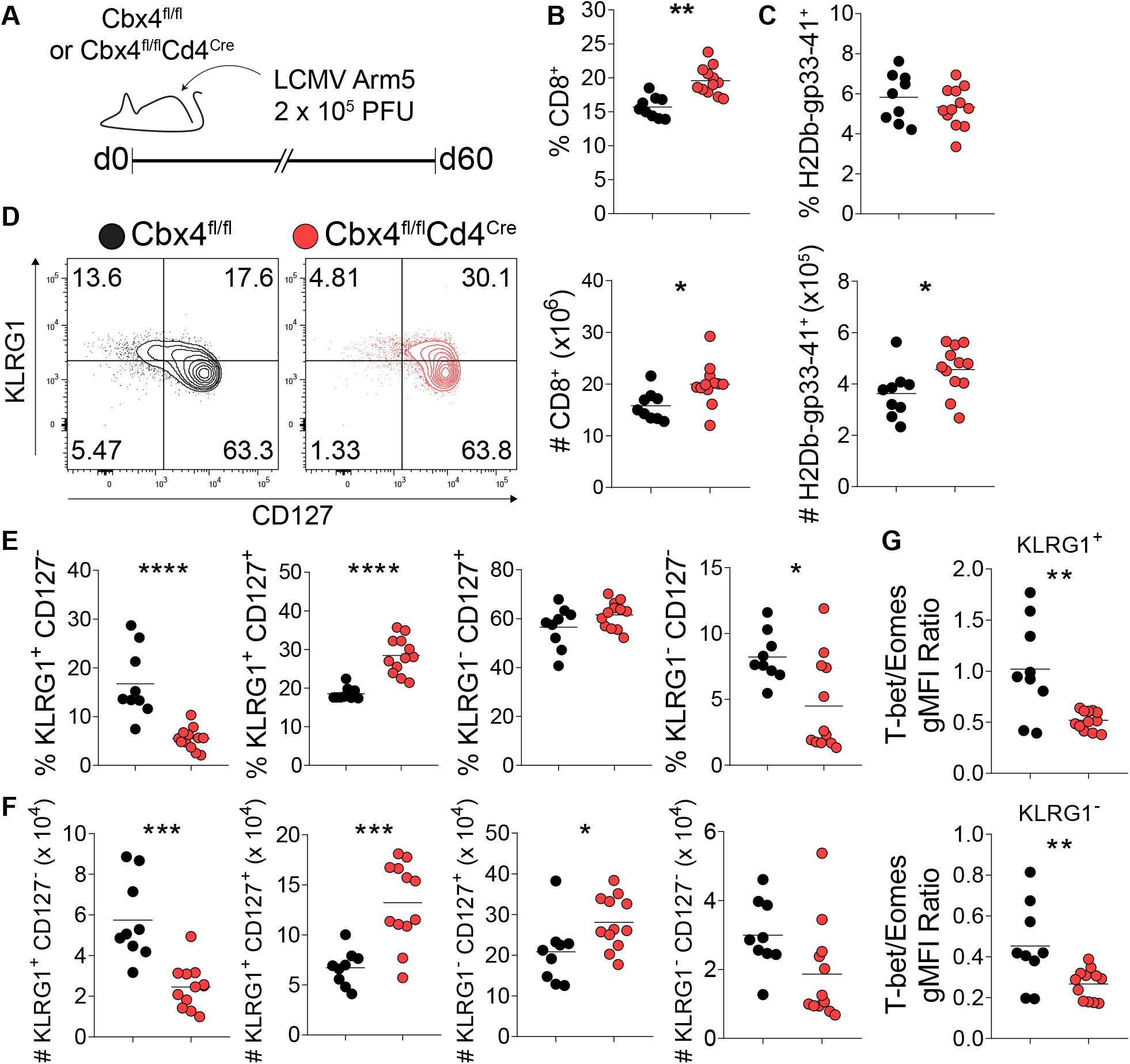
Cbx4 deficiency increases CD8 T cell memory formation. (A) Polyclonal Cbx4^fl/fl^ (black) or Cbx4^fl/fl^Cd4^Cre^ (red) mice were infected with acute infection LCMV and analyzed at 60 dpi. Frequency and total cell numbers of total CD8 T cell population (B) or LCMV-specific CD8 T cells (C) were measured by flow cytometry. The expression of KLRG1 and CD127 was analyzed by flow cytometry at 60 dpi (D), and frequency (E) and cell numbers (F) of each subpopulation were calculated. The expression of T-bet and Eomes was measured by gMFI in flow cytometry and T-bet/Eomes ratio was calculated using those values in both KLRG1^+^ and KLRG1^-^ populations (G). Representative contour plots for KLRG1 and CD127 expression are shown (D). Data are representative of two independent experiments (n ≥ 3). \**p* < 0.05, \*\**p* < 0.01, \*\*\**p* < 0.001, \*\*\*\**p* < 0.0001 by unpaired two-tailed Student *t* test (B, C, E-G).

To recapitulate this phenotype in an *in vitro* model system, we differentiate CD8 T cells to either effector- or memory-like cells using a previously established protocol (46). Consistently, we observed increased levels of the memory markers CD127 and CD62L in Cbx4-deficient cells, regardless of the polarized phenotypes (Suppl.Fig. 3). In summary, deficiency in Cbx4 leads to increased expression of memory-associated surface markers and transcription factors in CTLs, both *in vivo* and *in vitro*.

### Repression of memory phenotype by Cbx4 is primarily dependent on SIM1-2 domains

We next investigated the mechanism through which Cbx4 impacts CD8 T cell differentiation. As Cbx4 protein can mediate PcG-dependent repression and function in parallel as an E3 ligase enzyme (32, 37–39), we enforced the expression, in either wild-type or Cbx4-deficient P14 cells, of wild type or mutant Cbx4 cDNAs lacking key functional domains: (1) ΔChromo, with an amino acid substitution at chromodomain that prevents its binding to H3K27me3 residues; (2) ΔSIM1-2, that lacks both SUMO-interacting motifs (Fig. 3A).

**Figure 3.**
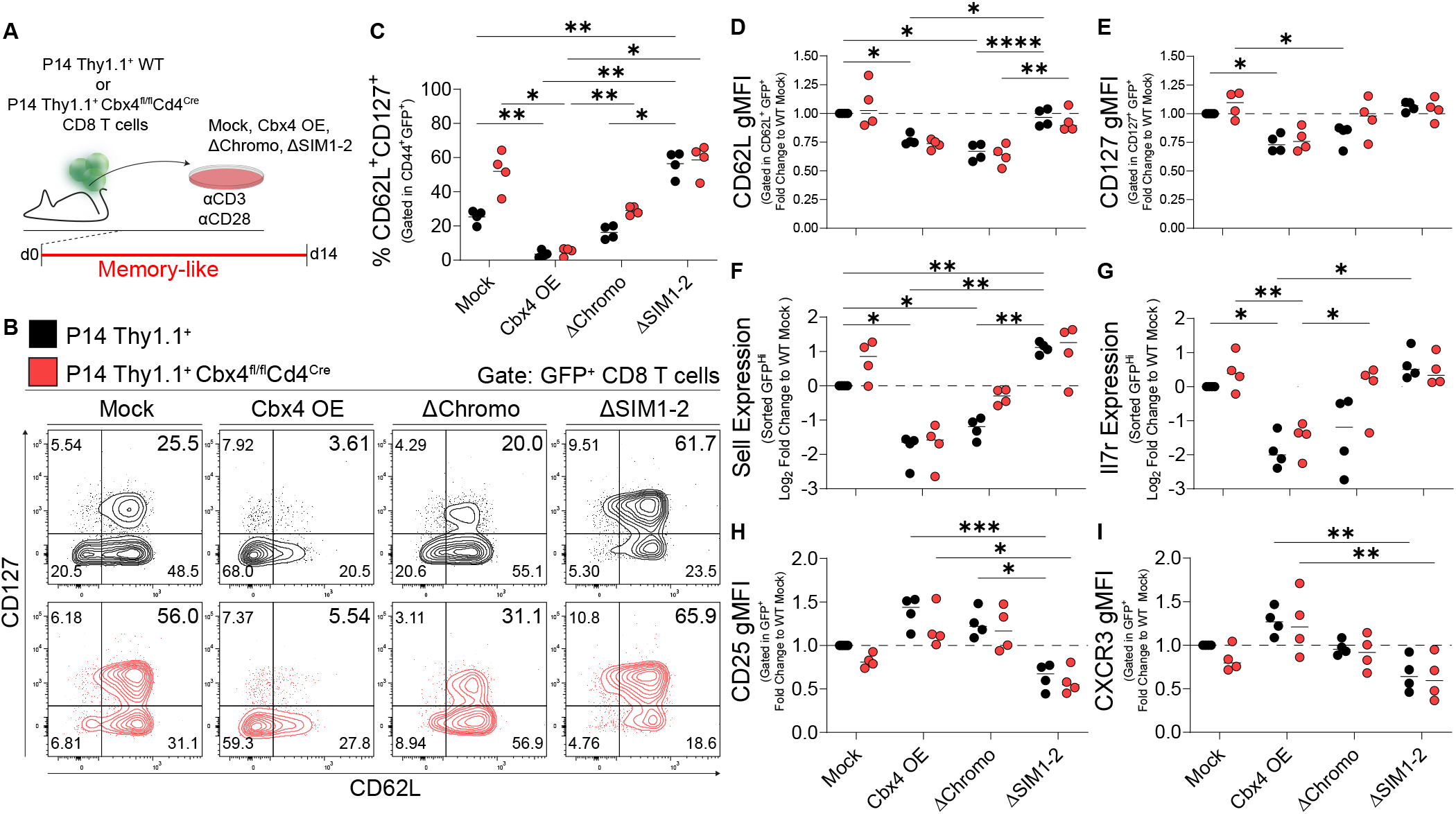
Differential requirement of Chromo and SIM1-2 Cbx4 domains for CTL differentiation. (A) *In vitro*-activated P14 Thy1.1^+^ WT (black) or P14 Thy1.1^+^ Cbx4^fl/fl^CD4^Cre^ (red) cells were transduced with either of Mock (control), wild type Cbx4 mRNA, Cbx4 ΔChromo mutant or Cbx4 ΔSIM1-2 mutant, and then polarized *in vitro* to memory-like phenotype (10U/mL of rhIL-2, 10 ng/mL of rhIL-7 and 10 ng/mL of rhIL-15) until day 14 of culture. The frequency of CD8 T cells expressing both CD62L and CD127 (C), as well as the expression of each marker measured by gMFI (D, E) was analyzed by flow cytometry. Gene expression for *Sell* (CD62L) and *Il7r* (CD127) was quantified by RT-qPCR using reporter GFP^+^-sorted cells (F, G). Expression of effector-related markers CD25 and CXCR3 (H, I). All gMFI and mRNA expression data are normalized to P14 Thy1.1^+^ WT transduced with Mock. Representative contour plots for CD62L and CD127 cytometry are shown (B). Data are representative of four (C-I) independent experiments. \**p* < 0.05, \*\**p* < 0.01, \*\*\**p* < 0.001, \*\*\*\**p* < 0.0001 by two-way ANOVA with Tukey’s test for multiple comparisons.

Cbx4 KO CD8 T cells activated and differentiated *in vitro* to a memory-like phenotype displayed accentuated expression of CD62L and CD127 during cell culture, and complementation of Cbx4-deficient cells with wildtype Cbx4 counteracted this phenotype (Fig. 3B, C). Overexpression of ΔChromo mutant reproduces to a lesser degree the effect seen upon wild-type isoform overexpression, indicating that the Cbx4 H3K27me3 binding function has a partial contribution to its role in the commitment to effector cell differentiation. On the other hand, deletion of SIM sequences not only reversed wild-type isoform Cbx4 overexpression impact but induced the opposite effect, promoting memory phenotype in both P14 WT and P14 Cbx4 T KO cells. Overexpression of ΔSIM1-2 mutant in P14 WT CD8 T cells raised the frequency of CD62L^+^CD127^+^ population to levels observed in Mock-transduced P14 Cbx4 KO cells, suggesting that overexpression of SIM-deficient Cbx4 isoform might be competing with endogenous Cbx4 function, and potentially acting as a dominant negative version of the protein (Fig.3.C). The same patterns were observed for the expression of memory-related markers CD62L and CD127 at protein (Fig.3.D, E) and transcriptional (Fig.3.F, G) levels. Expression of markers related to the effector phenotype, CD25, and recall response, CXCR3, corroborated these data, showing an increase of both proteins upon wild-type Cbx4 overexpression while ΔChromo mutant overexpression partially reproduced wild type Cbx4 effect and ΔSIM1-2 mutant reversed the effect (Fig.3.H, I).

The fact that we observed that both chromodomain and SIM domains are required for effector CD8 T cell differentiation is consistent with the fact that it has been shown in mouse embryonic fibroblast that conjugation of SUMO at the Cbx4 SIM domain is essential for recruitment of cPRC1 to H3K27me3 in genome *loci* and control of PRC1 repressive activity (54). Moreover, it was reported that Cbx4-mediated SUMOylation of Ezh2 promoted its recruitment and enhanced Ezh2 methyltransferase activity, demonstrating that Cbx4 regulates PRC2 (55). Therefore, our results indicate that Cbx4 SIM domains might regulate cPRC1 and PRC2-dependent repressive mechanisms in CD8 T cells. Further studies investigating Cbx4 SUMOylation and targets for Cbx4 SUMO E3 ligase function in CD8 T cells are needed to fully define its role in CD8 T cell differentiation. In addition, looking into the crosstalk between Cbx4 and other PcG proteins in CD8 T cells, especially Ezh2, and its potential interaction with H3K27me3 residues through the chromodomain is needed to understand how this protein controls Polycomb-mediated repressive mechanisms during CD8 T cell differentiation. Finally, the ability of Cbx4 to also bind to H3K9me3 adds another potential layer of complexity to the epigenetic regulation of T cells by this protein (37).

Taken together, our data demonstrates that Cbx4, a Polycomb-group protein, participates in the control of CD8 T cell differentiation. We observed that both Cbx4 SIM1-2 domains and chromodomain are required for the repression of memory phenotype, however SIM1-2 might play a more dominant role in this process. Further studies better characterizing the role of Cbx4 to CD8 T cell functionality and clarifying the mechanisms by which Cbx4 perform its role as regulator, identifying its targets and partners, will help clarify how the Polycomb-dependent complex molecular network regulates CD8 T cell differentiation.

