## Supplemental Figures for "Polycomb Repressive Complex 1 subunit Cbx4 positively regulates effector responses in CD8 T cells"

#### Supplementary Figures Legends

**Supplementary Figure 1. Even KLRG1$^{hi}$ Cbx4-deficient CD8 T cells skew to memory phenotype.** Relative expression of Cbx4 mRNA (quantified by RT-qPCR) of *in vitro*-activated C57BL/6 CD8 T cells transduced with RV expressing shCD4 or shCbx4 and polarized to effector-like phenotype (A). Frequency (B) and cell numbers (C) of total CD8, Thy1.1$^{+}$ or reporter Ametrine$^{+}$ population of P14 Thy1.1$^{+}$ cells transduced with control shRNA (shCD4, black) or shRNA targeting Cbx4 (shCbx4, red) and transferred to receptor mice in acute viral infection model (Figure 1). CD8 T cells knockdown for Cbx4 or control CD8 T cells were sorted based on KLRG1 expression (KLRG1$^{Hi}$) and the transcriptional profile analyzed by RNA-Seq. The volcano plot shows differentially expressed genes (Log2 Fold Change > 1 and adjusted p-value < 0.05) highlighted in red (D). Data are representative of ≥ three *in vitro* differentiation experiments (A) or two independent *in vivo* acute infection model experiments (n ≥ 3) (B, C and D). *$p < 0.05$, **$p < 0.01$, ***$p < 0.001$, ****$p < 0.0001$ by unpaired two-tailed Student *t*-test.

**Supplementary Figure 2. Cbx4 deficient CD8 T cells increase CXCR3-expressing populations.** Cbx4$^{fl/fl}$ or Cbx4$^{fl/fl}$Cd4$^{Cre}$ mice infected with acute infection LCMV were analyzed at 60 dpi. Frequency (A) and total cell numbers (B) of CD8 T cells subpopulations expressing KLRG1 and/or CXCR3 were measured by flow cytometry. Data are representative of two independent experiments (n ≥ 3). *$p < 0.05$, **$p < 0.01$, ***$p < 0.001$ by unpaired two-tailed Student *t*-test (A, B).

**Supplementary Figure 3. Cbx4 knockdown in *in vitro* differentiated CD8 T cells leads to upregulated expression of memory-associated surface markers.** *In vitro*-activated CD8 T cells were transduced with shCD4 or shCbx4, polarized to effector-like (200 U/mL of rmIL-2) or memory-like (20U/mL of rmIL-2, 10 ng/mL of rmIL-7 and 10 ng/mL of rmIL-15) phenotype and analyzed at day 6 or 14, respectively (A). The frequency of cells expressing memory-related surface markers CD127 (B) and CD62L (C) was measured by flow cytometry in both polarization conditions. Representative contour plots for CD62L and CD127 expression are shown (B and C, left). Data are representative of at least four *in vitro* polarization experiments. *$p < 0.05$, **$p < 0.01$, ***$p < 0.001$ by paired two-tailed Student *t*-test.

### Supplemental Figure 1

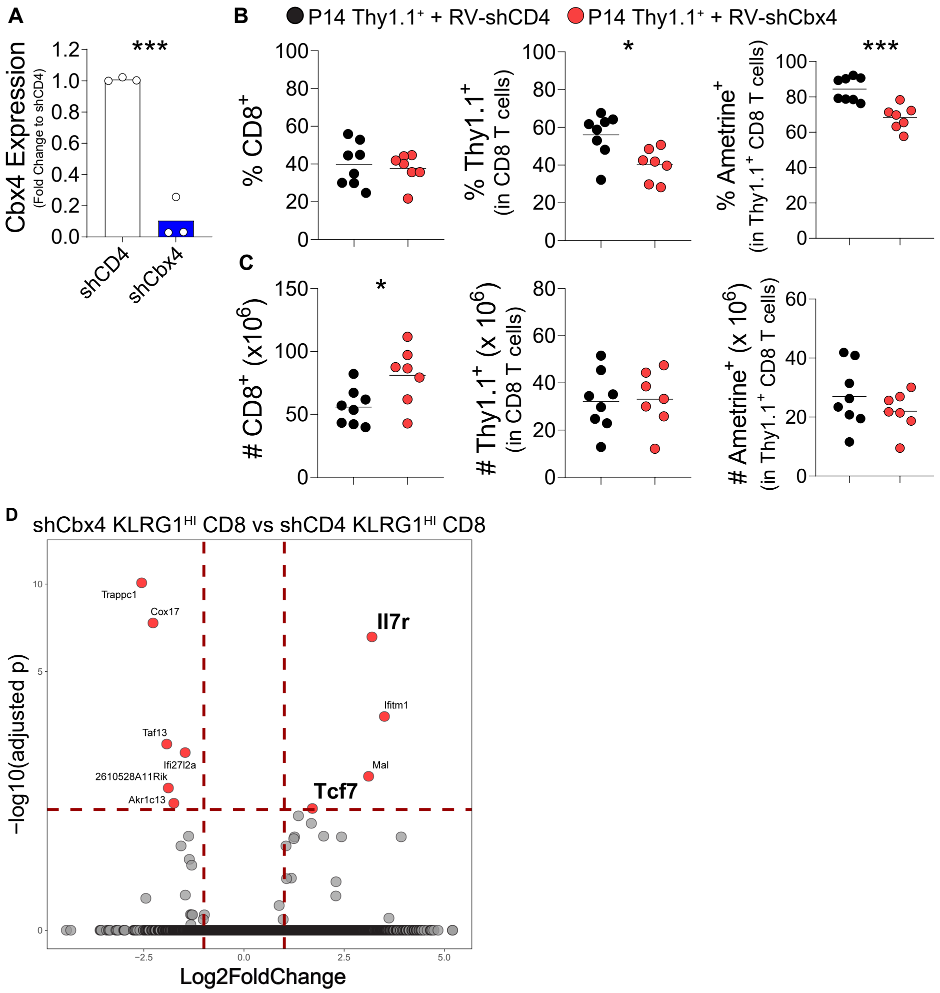

### Supplemental Figure 2

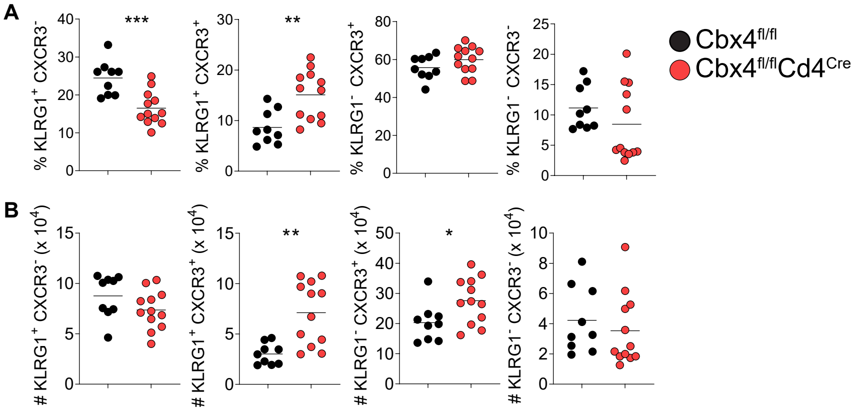

### Supplemental Figure 3

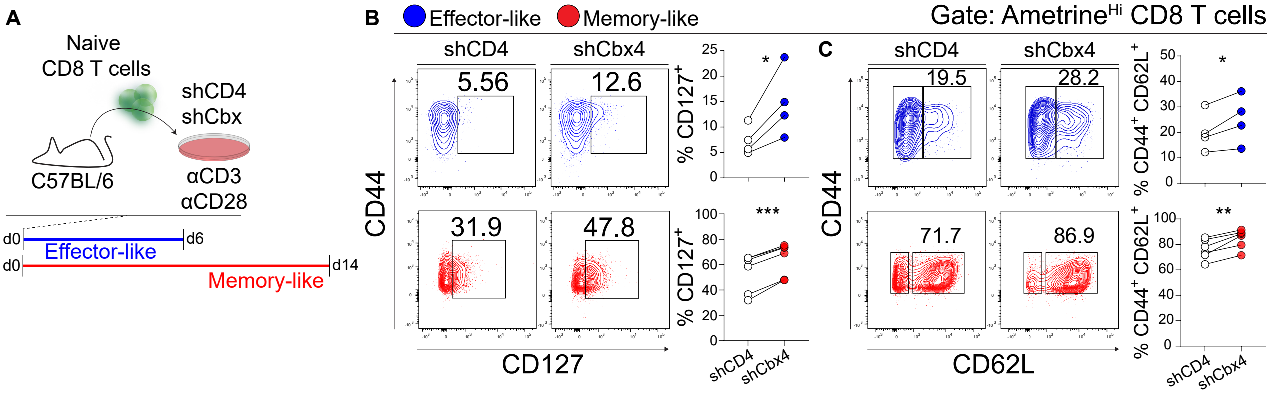
